# Linked-Read sequencing resolves complex structural variants

**DOI:** 10.1101/231662

**Authors:** Sarah Garcia, Stephen Williams, Andrew Wei Xu, Jill Herschleb, Patrick Marks, David Stafford, Deanna M. Church

## Abstract

Large genomic structural variants (>50bp) are important contributors to disease, yet they remain one of the most difficult types of variation to accurately ascertain, in part because they tend to cluster in duplicated and repetitive regions, but also because the various signals for these events can be challenging to detect with short reads. Clinically, aCGH and karyotype remain the most commonly used assays for genome-wide structural variant (SV) detection, though there is clear potential benefit to an NGS-based assay that accurately detects both SVs and single nucleotide variants. Linked-Read sequencing is a relatively simple, fast, and cost-effective method that is applicable to both genome and targeted assays. Linked-Reads are generated by performing haplotype-level dilution of long input DNA molecules into >1 million barcoded partitions, generating barcoded short reads within those partitions, and then performing short read sequencing in bulk. We performed 30x Linked-Read genome sequencing on a set of 23 samples with known balanced or unbalanced SVs. Twenty-seven of the 29 known events were detected and another event was called as a candidate. Sequence downsampling was performed on a subset to determine the lowest sequence depth required to detect variations. Copy-number variants can be called with as little as 1-2x sequencing depth (5-10Gb) while balanced events require on the order of 10x coverage for variant calls to be made, although specific signal is clearly present at 1-2x sequencing depth. In addition to detecting a full spectrum of variant types with a single test, Linked-Read sequencing provides base-level resolution of breakpoints, enabling complete resolution of even the most complex chromosomal rearrangements.

## Introduction

Structural variants (SVs), including balanced and unbalanced events, are known to be important contributors to a wide variety of human phenotypes including Charcot-Marie-Tooth syndrome and numerous intellectual disability syndromes [1,2]. Recent work has demonstrated that SVs are more prevalent in the general population than previously appreciated and that many structural events are more complex than previously thought [3,4]. Population-wide genome studies indicate that structural variation accounts for most of the differences between individual genomes and can explain differences in susceptibility to chromosomal rearrangements between groups [5–7]. But despite the widespread recognition of the contribution of SVs to health and disease, they remain poorly understood, in large part because they are generally undetectable with standard, short-read sequencing approaches that have enabled the majority of genome analyses.

Short-read sequencing is fundamentally limited by the lack of long-range information necessary to span many SVs or to map across repetitive regions of the genome, where SVs are known to cluster [6,8,9]. Presently, array comparative genome hybridization (aCGH) and karyotyping are the predominant screening tests used for genome wide assessment of large (>50bp) SVs [10–12]. Array CGH is often the first approach due to ease of workflow, resolution and ability to target known regions of interest [2,11]. However, aCGH is limited in that it can only detect unbalanced events and does not provide breakpoint resolution [11]. Conventional karyotyping is able to detect microscopically visible aneuploidies and large SVs with a lower limit of detection of ∼5-10Mb. Using fluorescent in situ hybridization probes (FISH), instead of g-banding, provides for increased resolution at targeted sites. While these two orthogonal approaches are complementary to next-generation sequencing based approaches, they increase the workflow and analysis burden. Recently, laboratories have demonstrated that low coverage long-insert WGS can identify both CNVs and balanced rearrangements at a cost and speed similar to aCGH/ karyotype, but the protocols can be difficult and time-consuming to establish [3,13].

None of the genomic screening assays currently available are capable of detecting the full spectrum of possible variants. Thus, when analyzing samples to understand the genetic underpinnings of disease, multiple assays often need to be employed to identify all potential variants [4,11]. Due to the additional cost and experimental burden involved in assaying SVs, they have been widely excluded from population-wide studies of genomic variation and next-generation-based clinical testing. As a result, when a study identifies a structural variant, it is often difficult to interpret whether the event is pathogenic or a neutral allelic variant present commonly in the population.

To address these variant detection shortcomings, we developed a technology that retains long-range information while maintaining the power, accuracy, and scalability of short-read sequencing. The 10x Genomics Chromium™ Genome solution utilizes haplotype-level dilution of high molecular weight DNA molecules into >1 million barcoded partitions to create a novel data type referred to as ‘Linked-Reads’. These Linked-Reads enable high-resolution genome analysis with minimal DNA input (~1 ng) [14]. We recently described both laboratory and analysis updates to our system, with a particular focus on the ability to identify small variants and to reconstruct long range haplotypes [15]. We have also described the ability of Linked-Reads to enable *de novo*, diploid assembly of individual genomes [16].

In the work described here, we performed 30x Linked-Read genome sequencing on a set of 23 samples with known balanced, unbalanced or complex SVs from either 1) the GetRm CNVPanel (unbalanced events) or 2) the Coriell general Cell Repository (balanced events). These cell lines have multiple, orthogonal assays confirming the presence of their described structural variants. Twenty-seven of the 29 structural variants assayed over 23 samples were identified as high quality variant calls, and one additional call was filtered to a secondary list of candidate structural variants as the breakpoints overlapped a segmental duplication, a feature known to complicate genome analysis. The remaining undetected event is a balanced translocation with a breakpoint in a heterochromatic region of chromosome 16. This region is represented by Ns in the reference assembly and will be invisible to any sequence-based method relying on the reference genome [17]. We also demonstrate that these events can be called with limited sequencing data. The Linked-Read approach, coupled with 30x sequencing, allows for the simultaneous identification of both small and large variation from a single library and analysis pipeline.

## Results

Our reference-based pipeline, Long Ranger, uses a set of novel algorithms to detect a full range of genomic variants using Linked-Read sequence data as input. The algorithms that support detection of large structural variants look for deviations in barcode coverage or unexpected barcode overlap. Barcode coverage, the number of molecules known or inferred to be present over a genomic region, is more even across the genome than read coverage and provides a modest improvement over read coverage in detecting CNVs at full depth sequencing (Fig 1a). Assessing unexpected barcode overlap allows for the detection of balanced events, and many unbalanced events, and is a method unique to Linked-Reads. In samples that match the reference genome, there should be little to no barcode overlap when the number of shared barcodes is plotted between two genomic regions separated by a genomic distance greater than the length of the input DNA (illustrated in Fig 1b). Each type of structural rearrangement results in a different pattern of barcode overlap that can reveal the type of variant as well as the coordinate(s) and orientation of breakpoints relative to each other. Examples of the canonical barcode overlap patterns for a deletion and an inversion are shown in figures 1 c and d, additional patterns are shown in supplemental figure 1. We assessed the performance of each of these barcode-based detection methods individually in our panel of samples covering a wide range of structural variant types (Table 1).

**Figure 1.**
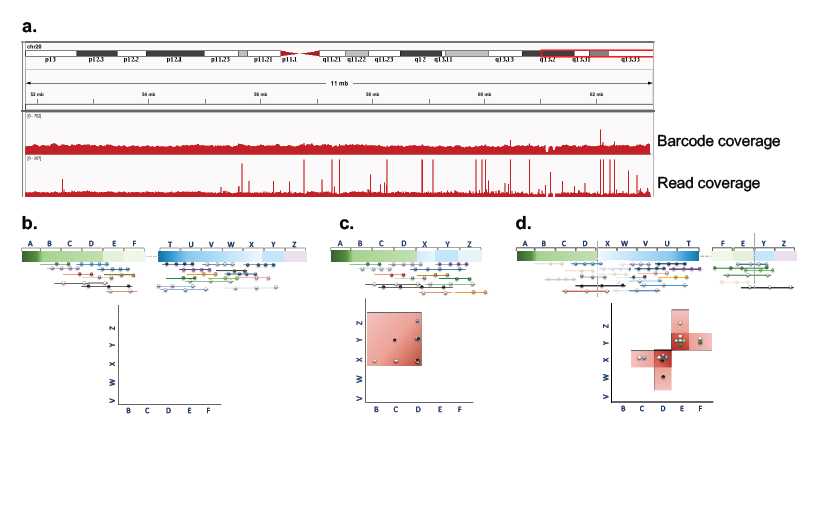
Barcode coverage (a) and barcode overlap (b-d) visualizations. **a**. Comparison of barcode coverage and read coverage over a 11Mb region of chr20 in NA12878. **b**. Schematic of two distant regions; there is no barcode overlap between these regions because they do not co-occur on the same long input DNA molecules and thus no signal can be visualized in the barcode overlap matrix plot. **c**. Schematic of a deletion that spans from segments E to W, bringing distant regions into proximity such that input molecules will now span both regions and barcode sharing will be seen between the distant regions. An anomalous overlap is seen in the matrix in a pattern that predicts a deletion. The highest degree of overlap occurs between segments D and X, which indicates the breakpoints of the deletion. **d.** Schematic of an inversion encompassing the region from segment E to X. The resulting anomalous barcode overlap pattern in the matrix is predictive of an inversion. The highest degree of overlap occurs between segments D/X and E/Y, corresponding to the inversion breakpoints.

**Table 1.** Long Ranger SV analysis of 23 Coriell samples with multiply-confirmed balanced or unbalanced SVs

|  |  | Samples | Event in Sample | Barcode Coverage | Barcode Overlap | Both Methods |
| --- | --- | --- | --- | --- | --- | --- |
| Copy Number Losses | Terminal Events | GM06936 | Deletion | Yes | No† | Yes |
|  |  | GM10989 | Deletion | Yes | No† | Yes |
|  |  | GM20027 | Aneuploidy | Yes | No† | Yes |
|  |  | GM21886 | Ring chromosome | Yes | No† | Yes |
|  |  | GM06226* | Derivative chromosome | Yes | No† | Yes |
|  |  | GM21699* | Derivative chromosome | Yes | No† | Yes |
|  |  | GM14485* | Derivative chromosome | Yes | No† | Yes |
|  | Non-Terminal Events | GM09888 | Deletion | Yes | Yes | Yes |
|  |  | GM14164 | Deletion | Yes | Yes | Yes |
|  |  | GM09216 | Deletion | Yes | Yes | Yes |
|  |  | GM10925‡ | Deletion | No | Yes | Yes |
| Copy Number Gains | Terminal Events | GM05966 | Duplication | Yes | No† | Yes |
|  |  | GM01416 | Aneuploidy | Yes | No† | Yes |
|  |  | GM05067 | Partial Aneuploidy | Yes | No† | Yes |
|  |  | GM16362 | Partial Aneuploidy | Yes | No† | Yes |
|  |  | GM20556 | Isodicentric chromosome | Yes | No† | Yes |
|  |  | GM06870 | Isodicentric chromosome | Yes | No† | Yes |
|  |  | GM06226* | Derivative chromosome | Yes | No† | Yes |
|  |  | GM21699* | Derivative chromosome | Yes | No† | Yes |
|  |  | GM14485* | Derivative chromosome | Yes | No† | Yes |
|  | **Non-Terminal Events** | GM09367 | Duplication | Yes | Yes | Yes |
| **Copy Neutral Events** | **Translocations** | GM06226* | Derivative chromosome | No† | Yes | Yes |
|  |  | GM21699* | Derivative chromosome | No† | Yes | Yes |
|  |  | GM14485* | Derivative chromosome | No† | Yes | Yes |
|  |  | GM22765 | Balanced translocation | No† | Yes | Yes |
|  |  | GM10207 | Balanced translocation | No† | Yes | Yes |
|  |  | GM18825 | Balanced translocation | No† | Candidate call | Candidate call |
|  |  | GM22709§ | Balanced translocation | No† | No | No |
|  | **Inversions** | GM21075 | Inversion | No† | Yes | Yes |

\*Sample contains multiple structural variants. †Algorithm not expected to detect this variant type. ‡Deletion in GM10925 falls in a segmental duplication; was called with high-quality score by Long Ranger but filtered as a likely false positive. §Balanced translocation in GM22709 falls within a heterochromatic region on chromosome 16 where there are known gaps in the reference assembly.

The 23 Coriell samples selected for this analysis represent a full range of structural variants including large deletions, duplications, inversions, balanced translocations and unbalanced translocations. Samples may contain multiple events (Table1). For example, a sample containing an unbalanced chromosome arrangement will have structural events representing a translocation, deletion, and duplication. Together, the barcode overlap and barcode coverage methods detected 28 of 29 total breakpoints, correctly characterizing 22 of the 23 samples tested (Table 1).

### Detecting Copy Number Variants

The copy number variants in our sample set included 6 large deletions, 4 large duplications and 4 large, unbalanced translocations (each with 1 large deletion and 1 large duplication) (supplemental table 1). We first sought to identify these variants using the barcode overlap method. Figure 2a shows a Loupe visualization of barcode overlap along chromosome 2 for sample GM09216, where Long Ranger correctly called a deletion allele. The matrix view reveals a pattern of barcode overlap signifying a deletion, and the linear view (top) reveals the positions of the breakpoint, which are calculated based on the maximum intensity of barcode overlap between regions. Also shown is an example of a duplication event detected on chromosome 6 in sample GM09367 (Fig 2b). At full depth sequencing (128Gb), the barcode overlap method detected all 5 of the non-terminal deletions and duplications in our sample set (Table1). As expected, because it requires the ability to examine barcode sharing patterns on both sides of a breakpoint, this method did not detect terminal deletions or duplications.

**Figure 2.**
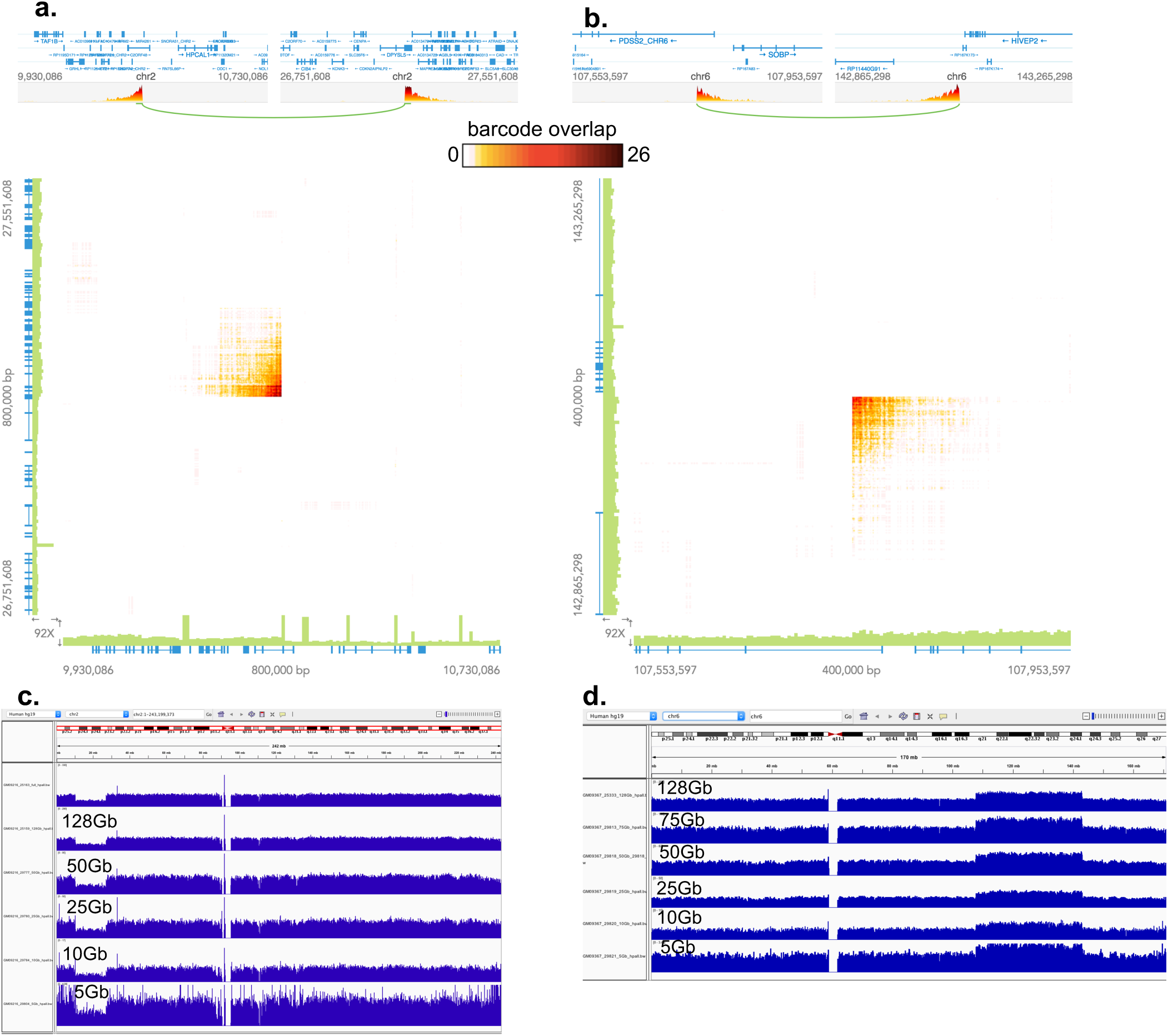
Copy number variants detected with barcode overlap and barcode coverage. Visualizations of a deletion event in sample GM09261: 46,XY,del(2)(p25.1p23) **(a. and c.)** and a duplication event in sample GM09367: 46,XX,dup(6)(q21q24) **(b. and d.)**. **a and b** Barcode overlap linear (top) and matrix (bottom) views of these events with 128Gb sequence. These events were not called by barcode overlap at lower sequence depths. **c. and d.** IGV tracks showing barcode coverage in the event regions with sequence depths of 128Gb down to 5Gb, as indicated. Both events were called by the barcode coverage method at all sequence depths tested.

We next examined the performance of the barcode coverage algorithms for calling CNVs. Deviations in barcode coverage that signal a copy number variant are visible in the barcode coverage tracks, as shown for two samples in figure 2c-d. The first example shows sample GM09261, the same deletion variant that was called with barcode overlap in figure 2a. Here, the large deletion in the long arm of chromosome 2 is evident by a reduction of barcode coverage in the 2p region (fig 2c). Similarly, the duplication on the short arm of chromosome 6 in sample GM09367 is evident as an increase of barcode coverage in the 6q region (fig 2d). The barcode coverage algorithm was able to detect 19 out of 20 known CNVs, including all 9 of the terminal events (Table 1). The undetected variant, a deletion in sample GM10925, falls within a segmental duplication and was filtered by the algorithm as a likely false positive (Table1). However, this event was detected with the barcode overlap method, demonstrating the advantage of utilizing multiple detection methods for variant calling.

Downsampling of sequence data was performed *in silico* to determine the minimum sequence depth required to detect each CNV. Due to the evenness provided by barcode coverage, the deletion and duplication signals for these two samples are detectable even with coverage as low as 5Gb (~1x genomic read coverage) (Fig 2 c and d). The barcode overlap method was less robust with reduced sequencing and did not call CNVs with less than 50Gb of sequence. This result is expected, given that the algorithm was designed for use with full-depth data. However, there was an observable signal in the barcode overlap data with as little as 5-10Gb for many of the samples, indicating that the algorithm is likely extensible to lower depth data. (supplemental table 2)

### Detecting Balanced Structural Variants with Barcode Overlap

Because the barcode overlap algorithm does not rely on coverage differences for variant detection, this method has the power to detect balanced SVs including translocations and inversions from Linked-Read data. Figure 3a shows the Loupe Matrix view for sample GM21075 (sequenced to 128Gb) where Long Ranger correctly called a balanced inversion on the long arm of chromosome 9. Seven of the 8 balanced events in our dataset were called accurately with the barcode overlap method at full depth. *In silico* sequence downsampling revealed that Long Ranger could call these balanced events with as little as 10x read coverage (50Gb), with specific signal present down to 1-2x read coverage (5-10Gb) (Fig 3b). The undetected event in sample GM22709 is a balanced translocation on the long arm of chromosome 16, where there is a known gap in the reference assembly.

**Figure 3.**
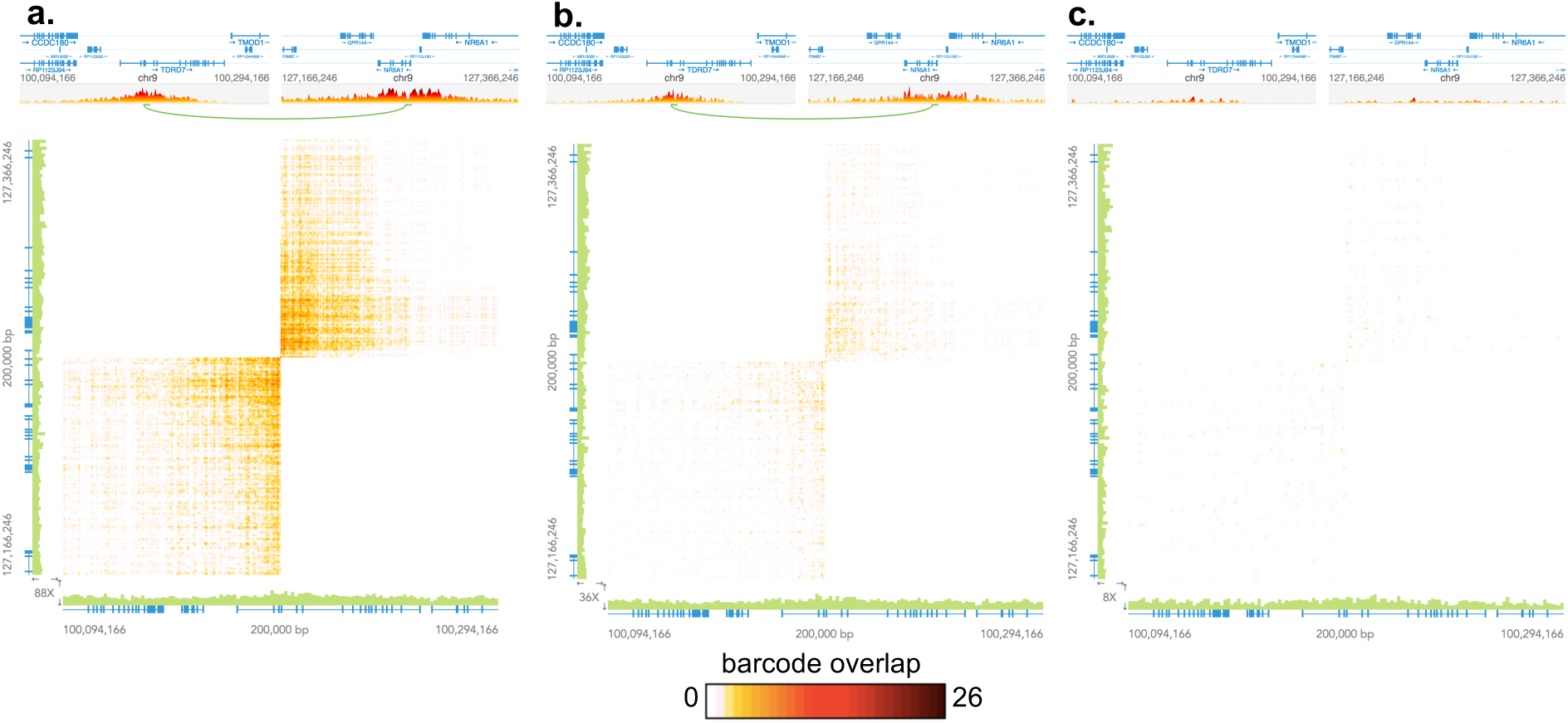
Detection of event for GM21075: 46,XY,inv(9)(q22.3q34.1). **a.** Barcode Matrix view showing a balanced inversion detected on the long arm of chromosome 9 with 128Gb of Linked-Read sequence data. **b.** Barcode matrix view of the same inversion event shown with only 50Gb of Linked-Read sequence coverage, the lowest coverage at which Long Ranger called this event. **c.** The same event shown with 10Gb of Lined-Read sequence coverage showing that there is signal for this event in the data at this coverage level, even though Long Ranger does not make the definitive call.

### Complex event resolution

The Long Ranger pipeline calls each breakpoint individually and does not integrate multiple related breakpoints into single complex structural events. As a result, the samples in our set that harbor derivative, unbalanced chromosomes are not annotated as such. Instead, they have three separate structural variant calls comprising the event: translocation, deletion, and duplication (Table 1 asterisks). To fully understand the nature of the these complex genomic rearrangements, we manually examined each variant in the Loupe browser. Figure 4 shows the manual examination of an unbalanced translocation involving chromosomes 1 and 16. The Loupe linear view (4a top) shows the gene models near the breakpoints. The three events called with Long Ranger (translocation, terminal deletion and terminal duplication) are visualized in panels a,b and c, respectively. However, the uneven barcode coverage along chromosome 16 indicates something more complicated than a simple duplication at this breakpoint (Fig. 4c).

**Figure 4.**
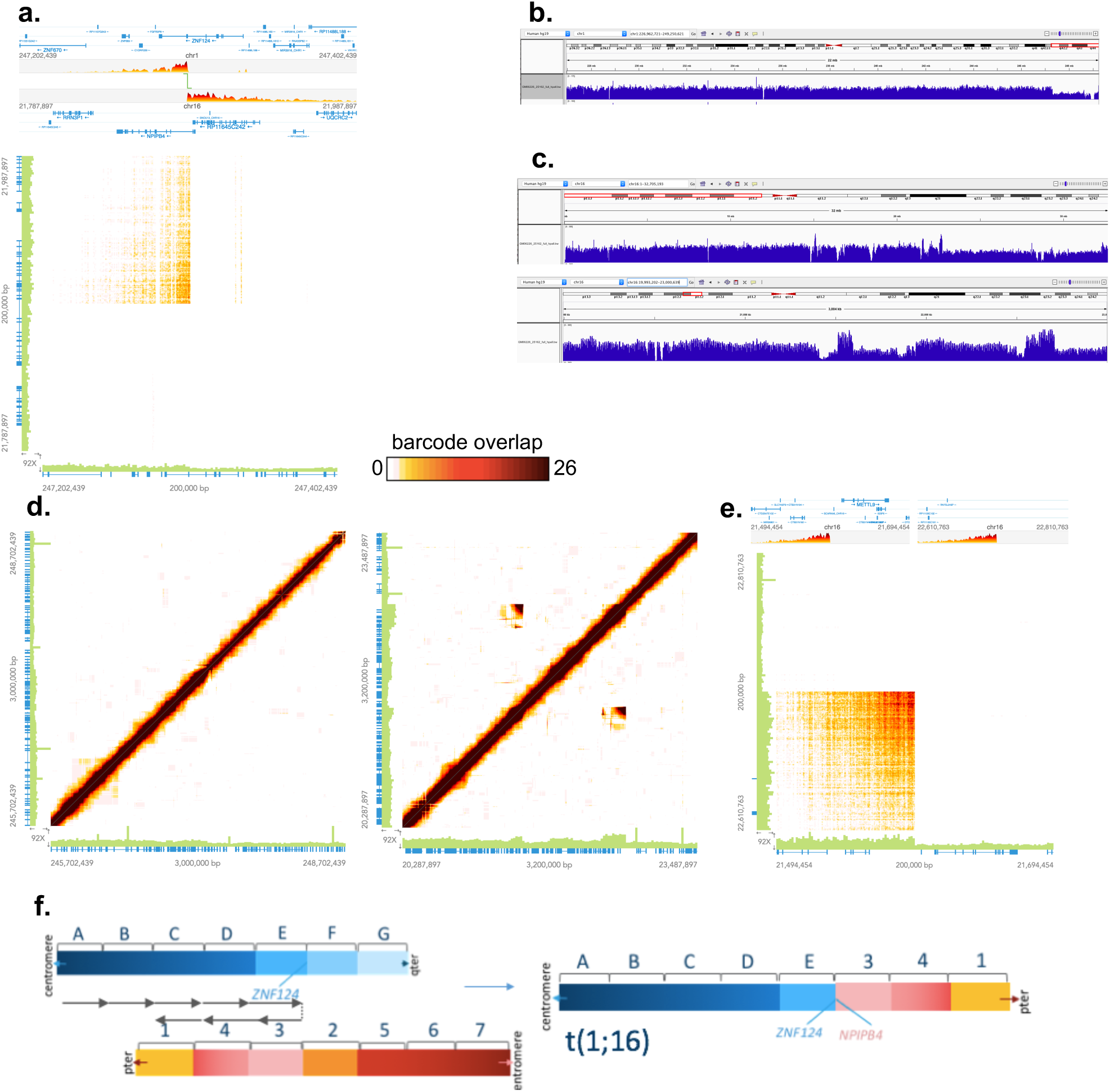
Derivation of a complex structural event in sample GM06226: 46,XY,der(1)t(1;16)(q44;p12) **a.** Barcode overlap linear (top) and matrix (bottom) views of the t(1;16) event. **b**. Barcode coverage view of the q arm of chr1 where a terminal deletion was called. **c**. Barcode coverage view of the p arm of chr16 where Long Ranger called a terminal duplication (top). A zoomed barcode coverage view near the breakpoint on chr16p shows barcode coverage variability (bottom). **d**. Barcode overlap signatures in regions of predicted breakpoints reveal an additional structural on chr16, seen as unexpected overlap above and below the diagonal. **e.** A zoomed view of one of these regions reveals an overlap pattern indicative of a translocation. **f**. A model for the structural rearrangement depicting a translocation on the background of an inverted chr16. This configuration explains the discontiguous duplication signal on chr16 observed in c.

To further resolve this event, we examined the patterns of barcode overlap revealed over the regions with the putative breakpoints with immediately adjacent regions-effectively plotting barcode sharing between a large region and itself (Figure 4d and e). In the self/self plot of chromosome 1, the diagonal line of barcode overlap indicates that this region does not have any structural variation bringing two distant regions into close proximity as the greatest amount of barcode overlap is always observed between a region and its immediately adjacent window. However, the decreased intensity in the heat map at the right/upper end of the diagonal indicates a depletion of shared barcodes in this region, consistent with the chromosome 1 deletion called with barcode coverage (Fig 4d, left). In the alignment of chromosome 16 against itself, unexpected barcode sharing is observed as signal above and below the diagonal, indicating rearrangement (Fig 4d, right). Furthermore, analysis of the barcode coverage pattern in this region reveals a pattern not consistent with a simple duplication as the increased barcode overlap is observed as two disparate signals within the region (Fig 4c). A closer look at this overlap shows a pattern that is consistent with an inversion (Fig 4e), however, this unexpected barcode overlap does not result in a structural variant call because >50% of the structural variant overlaps a region on the 10x Genomics Blacklist. This blacklist defines gaps and other ambiguous regions of the reference genome that have been found to raise spurious large-scale SV calls. This region is known to contain a rare inversion in the reference assembly, so the majority of samples processed will have barcode evidence for an inversion relative to the reference. Therefore, the translocation observed in this sample occurred on a chromosome 16 with an inversion relative to the reference assembly-effectively separating the duplication signal observed when mapped back to the reference (regions 1,3, and 4 in Fig 4f). A model consistent with all of the variations and anomalous overlap signals detected in this sample involves a translocation on the background of an inverted chromosome 16 (Fig 4f).

## Discussion

We have demonstrated the power of Linked-Reads to detect both balanced and unbalanced structural variants using only ~1ng of input DNA, a single sequencing library, and a single analysis pipeline (Long Ranger), thus presenting a testing modality that can resolve multiple variant types with a lower workload and cost. However, there are some limitations to the Linked-Read data type and the results in our study. Although Linked-Reads can resolve many repetitive elements and genome regions, highly repetitive sequences that are larger than the length of input DNA are not resolvable by Linked-Reads. This limitation is common to all technologies currently available, including long-read sequencing. An additional limitation in this study is the reliance on a reference sample for calling variants, which creates reference bias and the inability to call variants in regions that are not resolved in the reference, as was the case with the variant in the pericentric region on chromosome 16. To bypass any reference bias, Linked-Read data can also be used to perform diploid *de novo* assembly using our Supernova™ pipelines [16]. However, *de novo* assembly requires ~56x sequence coverage and is not feasible for all samples or projects.

A notable finding in this analysis of Long Ranger is that Linked-Reads added resolution not previously achieved for some of the more complex events and resolved breakpoints down to the base pair level. This additional complexity is consistent with other recent studies showing that complex structural events sometimes result from catastrophic chromosomal rearrangements involving dozens of breakpoints [3,18]. In the latter study, Linked-Reads were used to fully resolve three highly complex variants that could not be fully resolved by other methods employed, including long-insert whole genome sequencing, PCR or Sanger sequencing [3].

In a recent study reported in *Genome Medicine*, researchers used Linked-Read sequencing to characterize variations in metastatic tumors [19]. Employing novel bioinformatic analysis methods designed for the Linked-Read datatype, they detected tight clusters of SV breakpoints in a genomic region encompassing the known cancer driving gene, *FGFR2*. Local assembly of the Linked-Reads that spanned this region revealed that the specific breakpoints were unique between surgically excised samples from two different metastatic sites [19]. These results demonstrate the ability of Linked-Reads to detect somatic genome rearrangements at the sequence level.

The results presented here demonstrate the potential of Linked-Reads to replace two standard assays, aCGH and karyotype analysis, with a single assay. In another recently submitted study, we demonstrate the power of Long Ranger and Linked-Reads to detect single nucleotide variants and indels; performance metrics for these small variants matched those achieved with standard Illumina TruSeq libraries [15]. Thus, by exploiting the multiple layers of barcode signals that can be detected from the Linked-Read sequence data, the Long Ranger sequence analysis pipeline detects SNVs, indels, CNVs, and balanced SVs in a single workflow.

## Conclusions

Linked-Reads have the potential to provide just one test for resolving a full range of variants that currently require multiple testing modalities to target completely. All of the variants described herein, including SNVs, are automatic products of our open source analysis pipeline and are delivered in file formats compatible with most open source analysis tools, including Loupe. Our algorithms accurately called all but one of the variants tested, and we also see signal in the data even when the pipeline does not make a definitive call. Thus, there is still potential for more discovery in the data with algorithm improvements and further validation with different clinical samples.

## Methods

### Samples and DNA isolation

Cell lines were obtained from the NIGMS Human Genetic Cell Repository at the Coriell Institute for Medical Research (repository ID numbers are listed in Table s1). Frozen cell pellets were thawed rapidly at 37°C in 1ML PBS. High molecular weight DNA was then extracted following recommended protocols (https://assets.contentful.com/an68im79xiti/1Jw6vQfW1GOGuO0AsS2gM8/9dd2a091194693dd9dc11a1cd7768c17/CG00043GenomeReagentKitv2UserGuideRevA.pdf) and quanitfied on a Qubit^®^ Fluorometer system.

### Library construction

1.25 ng of high molecular weight DNA was loaded onto a Chromium™ controller chip, along with 10x Chromium™ reagents and gel beads following recommended protocols (https://assets.contentful.com/an68im79xiti/4z5JA3C67KOyCE2ucacCM6/d05ce5fa3dc4282f3da5ae7296f2645b/CG00022_GenomeReagentKitUserGuide_RevC.pdf). The initial part of the library construction takes place within droplets containing beads with unique barcodes (called GEMs). The library construction incorporates a unique barcode that is adjacent to read one. All molecules within a GEM get tagged with the same barcode, but because of the limiting dilution of the genome (roughly 300 haploid genome equivalents) the chances that two molecules from the same region of the genome are partitioned in the same GEM is very small. Thus, the barcodes can be used to statistically associate short reads with their source long molecule.

### Sequencing and analysis

Libraries were sequenced on a combination of Illumina^®^ HiSeq^®^ 4000 and HiSeq^®^ 2500 sequencers. Base calling and quality (BCL) files were used as input to the Long Ranger pipeline using default parameters (https://support.10xgenomics.com/genome-exome/software/pipelines/latest/using/wgs).

### Structural variation analysis

All Long Ranger runs were performed with a pre-release build of Long Ranger version 2.2. Two algorithms were used to call large structural variants. An algorithm assessing for unexpected barcode overlap between distant regions is included in Long Ranger v1.2. The second algorithm, which assesses for deviations of barcode coverage for detecting copy number events, is included with Long Ranger 2.2. 10x plans to release and open-source Long Ranger 2.2 in February 2018. Structural variants were manually reviewed, using the available karyotype and array CGH results available from Coriell to guide analysis to specific regions where variation was known to be present.

## Acknowledgements/contributions

High molecular weight DNA extractions and library preparations were carried out by Heather Ordonez and Cassandra Jabara. Kariena Dill provided scientific writing support and and prepared figures for submission.

---

**Figure S1. Theoretical patterns of barcode overlap a.** Schematic of a duplication event that spans from segments T to X, bringing distant regions into proximity such that input molecules span both regions and barcode sharing will be seen between regions that are distant in the reference. The anomalous overlap pattern illustrated in the matrix predicts a duplication. The highest degree of overlap occurs between segments E and X, which indicates the breakpoints of the deletion. **b.** Schematic of an inverted translocation of the region encompassing segments T to X. The resulting anomalous barcode overlap pattern in the matrix is predictive of a translocation. The highest degree of overlap occurs between segments D/X and E/Y, corresponding to the breakpoints. **c.** In samples with sequence matching the reference genome, barcode overlap between a region and itself will follow a linear pattern. Overlap is strongest along the diagonal, with decreasing overlap occurring as a function of distance.

